# How Acting Jointly Differs from Acting Side-by-Side: A Dual EEG Study

**DOI:** 10.1101/2025.04.18.649548

**Authors:** Martina Fanghella, Elena Mussini, Agnese Zazio, Francesca Genovese, Eleonora Satta, Guido Barchiesi, Alexandra Battaglia-Mayer, Marta Bortoletto, Corrado Sinigaglia

## Abstract

The distinction between acting jointly and acting side by side permeates our daily lives and is crucial for understanding the evolution and development of human sociality. While acting in parallel involves agents pursuing *individual* goals, acting jointly requires them to share a *collective goal*. Here, we used a dual electroencephalography (EEG) approach to explore the neural dynamics underlying joint and parallel action preparation. We recorded event-related potentials (ERPs) from 20 dyads while they had to transport an object in a video game, either jointly or in parallel, or individually. Both conditions were carefully matched for coordination demands and performance complexity, as confirmed by equal success rates. Our results revealed a distinctive pattern swap in ERPs during action preparation. In the early preparation phase, ERPs showed significantly higher amplitude during joint action than parallel action. This pattern reversed in the late preparation phase, with significantly reduced ERP amplitude in the joint compared to parallel action. Notably, the decrease in late ERPs correlated with higher reaction time (RT) variability in partners but not with participants’ own RT variability. The dynamic swap in neural activity suggests that different cognitive processes operate at distinct stages of action preparation. While initially sharing a collective goal may impose cognitive costs (reflected in higher early ERPs), this is offset by facilitated late action preparation, likely due to enhanced predictability of partners’ actions.

**Significance Statement:** Our study reveals distinct neural signatures differentiating joint from side-by-side actions. Through dual EEG recordings of twenty dyads performing complexity-matched tasks, we identified a distinctive “swap” in event-related potentials during action preparation. Joint actions initially showed higher early-phase amplitudes but significantly reduced late-phase amplitudes compared to parallel actions. Notably, this late-phase reduction correlated specifically with the variability of partners’ behavior. This suggests that sharing collective goals initially requires cognitive resources but facilitates action preparation through enhanced predictability of partners. These findings provide a neural framework for understanding the distinction between acting jointly and acting in parallel —a distinction that pervades our daily experiences and is crucial for understanding the development and evolution of human sociality.

## INTRODUCTION

Anyone who has ever danced a tango or cooked a hollandaise sauce knows that acting jointly differs from merely acting side-by-side. This distinction pervades our daily experiences (Gilbert 1990; Bratman, 1993) and marks a pivotal step in human social evolution (Tomasello et al., 2005) and development (Carpenter, 2009; Brownell, 2013).

Over the last two decades, research has extensively investigated the processes underlying joint action (Sebanz & Knoblich, 2021). Studies consistently show that agents represent others’ actions (Loehr et al., 2013; Kourtis et al., 2014, 2019; Meyer et al., 2013; Novembre et al., 2014; Satta et al., 2017; Sacheli et al., 2018) and synchronize (d’Ausilio et al., 2015; Repp & Su, 2013; Keller et al., 2014; Pezzulo et al., 2017; Bigand et al., 2024) during shared activities like clinking glasses, playing music, or dancing. Similar results have been obtained in monkeys (Ferrari-Toniolo et al., 2019; Lacal et al., 2022; Pezzulo et al., 2022). Moreover, there is evidence that sharing task representations affects joint action performance (Schmitz et al., 2017, 2018). Other studies point to temporal adaptation, the speeding up or slowing down of individual actions to match observed behavior (Konvalinka et al., 2010; Nowicki et al., 2013; Lelonkiewicz & Gambi, 2017).

However, others’ action representation and synchronization also occur when acting individually. People often represent others’ actions even when it impairs their performance (Atmaca et al., 2008; Atmaca et al., 2011), and agents share task representation also when they are not required to act jointly (Sebanz et al., 2003). While synchronization can facilitate interpersonal coordination (Nessler & Gilliland, 2009; Richardson et al., 2009), it can occur unintentionally (Varlet et al., 2015) and persist despite attempts to avoid it (Issartel et al., 2007; Ultzen et al., 2008).

All these findings indicate that the distinction between joint and parallel action remains underinvestigated. Our paper aimed to bridge this gap. We used dual electroencephalography (EEG) to record brain activity from two individuals simultaneously, exploring the neural dynamics that distinguish joint from parallel action. Joint action is minimally characterized by a *collective goal* rather than the *individual goals* driving parallel action (Searle, 1990; Gilbert, 2013; Bratman, 2014). Instead of simply comparing solo and dyadic actions, as previous studies have done (but see della Gatta et al., 2017; Barchiesi et al., 2022; Formica & Brass, 2024), we contrasted two-agent actions directed toward either collective or individual goals, while matching the tasks for movement and coordination complexity.

Participants in dyads played a video game in which they controlled cursors to navigate a path and transport shovels into an igloo. In the Joint Action condition, two players moved one shovel *together*; in the Parallel condition, each player transported their shovel *individually* alongside the other. In both conditions, players should monitor their partner’s cursor movements. In the Joint condition, the action’s outcome depended on the successful performance of both players. In contrast, in the Parallel condition, each agent pursued their individual goal independently while avoiding collisions with the other.

We focused on the behavioral and electrophysiological differences between joint and parallel action preparation. At the behavioral level, we considered success rate to evaluate task complexity across conditions and analyzed mutual alignment within participant dyads by measuring the association between their reaction times and reaction time variability. At the electrophysiological level, we used a cluster-based permutation analysis to identify differences in event-related potential (ERP) amplitude between the Joint and Parallel conditions across all electrodes during the action preparation phase. Finally, we bridged the behavioral and electrophysiological levels by exploring whether differences in ERP amplitude across conditions correlated with changes in self- and partner-related reaction times and their variability.

If sharing a collective goal provides a core definition of joint action, it should manifest in different behavioral patterns and neural dynamics compared to parallel action. Sharing a collective goal rather than pursuing individual goals should impact on the various stages of action preparation, even when the tasks match in complexity.

## MATERIAL AND METHODS

### Participants

Twenty dyads of right-handed participants (n = 40, 19 females, mean age 24.1) were recruited for the experiment. All participants had normal or corrected-to-normal vision and no history of psychiatric or neurological disorders. The research protocol was approved by the Local Ethics Committee and was carried out in accordance with the principles of the revised Declaration of Helsinki (World Medical Association General Assembly, 2008). Written informed consent was obtained from all the participants. To calculate the sample size, we ran a power analysis based on the modulations of contingent negative variation in a joint task reported by Kourtis et al. (2019). Although there are consistent differences in the data analysis, their task provides the most similar paradigm. This analysis was carried out with MorePower and showed that 16 dyads, i.e., 32 participants, were sufficient to replicate the effect they observed (Experimental design: 2×2×2 repeated measures ANOVA; effect of interest: 2 (joint/no joint); Power: 0.80; Partial Eta^2^: .219; calculated sample size: 32 participants).

### Experimental setting

Participants were paired in dyads and seated at desks in an electrically shielded chamber (a Faraday cage), 150 cm apart. Each participant faced their computer screen (ASUS VG248QE, 61 cm × 34.4 cm, 60 Hz refresh rate, 1920 × 1080 resolution) positioned 50 cm away. The task was a cartoon-like video game, adapted from Satta et al. (2017), programmed in Psychtoolbox-3 on MATLAB R2023b (The MathWorks Inc., Natick, MA, USA). A single PC drove both displays, and each participant controlled their actions using a Saitek X65 F Control System isometric joystick with their right hand.

EEG was recorded simultaneously from both participants using a g.HIamp amplifier (g.tec medical engineering GmbH, Schiedlberg, Austria). Data were acquired at 1200 Hz using two 64-electrode caps following the 10-20 system. The reference electrode was placed at FPz, and the ground electrode was placed at the nose tip.

### Task

Participants controlled cursor characters (puppets) with a height of 3 degrees of visual angle (DVA), using joystick force pulses to move from the center in different directions. Characters were identical, except for their clothing color: red for the left participant and green for the right. The task required participants to navigate their characters, grab a shovel, and transport it into an igloo along a path. The experiment included two conditions: in *Joint Action* (JA), participants jointly transported a single shovel, while in *Parallel Action* (PA), each participant independently transported their own shovel alongside the other.

Each trial started with a black fixation cross on a white background, lasting 2000 ms. Participants were then presented with the red and green characters and a central circle. They were asked to position their character inside the circle and hold their cursor still to maintain the character within it for a variable time interval (center holding time, CHT), lasting from 1000 to 2000 ms in 100 ms increments. Any cursor displacement outside the target resulted in an error and consequent abortion of the trial. Then, an instruction *c*ue was presented for 1500 ms, corresponding to the action preparation phase, followed by a go-signal, upon which participants moved as fast and accurately as possible.

In the JA condition, the cue was a white snow shovel positioned at the beginning of a ground path extending from the center to a white igloo (9 × 5 DVA), appearing as a peripheral target at one of four random positions (11 DVA eccentricity from the screen center). The go-signal consisted of the shovel changing color from white to red and green. Both participants then moved to grab and transport the shovel *jointly* toward the igloo, maintaining coordination by staying within the marked path and keeping their inter-cursor distance under 150 pixels. Success triggered a snow removal animation and changed the igloo’s color to red and green. The trial ended unsuccessfully, leaving the igloo white if either participant anticipated the go signal, left the path, or exceeded the allowed distance from their partner. The trial outcome depended on the joint performance of both participants.

The PA condition followed a similar structure but with two key differences: (1) two shovels appeared instead of one, and (2) participants acted independently. At cue onset, two white shovels appeared on the path. At the go-signal, these shovels changed color (left to red, right to green), matching each participant’s character color. Participants were pre-informed about shovel positions and color changes, enabling movement planning during the preparation phase. After the go-signal, each participant had to reach their shovels and *individually* transport them to the igloo without leaving the path. To prevent a collision, participants followed priority rules: the red player had priority in the left half of the path, and the green player in the right half. Overlapping positions (within 50 pixels) in the same path half resulted in trial failure for the player without priority. Unlike in JA, individual errors (anticipating the go-signal, leaving the path, violating priority rules) affected only the participant who made them. Participants were carefully instructed that their actions were side by side, not competitive (i.e., the order of arrival at the igloo was irrelevant) or cooperative (i.e., avoiding systematic taking turns in action execution to prevent collisions). Success was also tracked individually: the igloo changed color partially (red or green) for each successful participant, or entirely for concurrent success, and remained white if both participants failed. For trial structure visualization in both conditions, see Figure 1 B.

**Figure 1.**
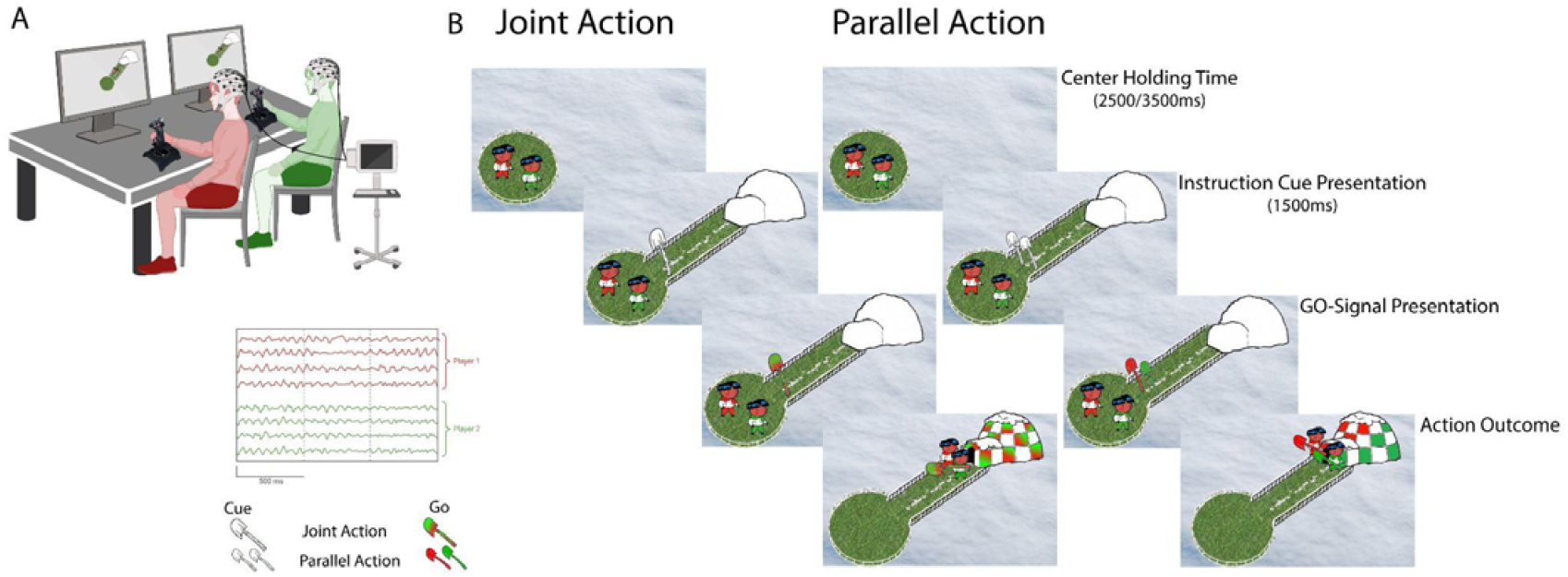
A. Experimental Setup: During the task, the two participants controlled a red character (on the left) or a green character (on the right) on their screens, using isometric joysticks, while their EEG activity was recorded simultaneously from the same amplifier. The EEG activity of interest consisted of a 1500 ms time window associated with the action preparation phase. B. Structure of a trial: JA and PA were matched for visual stimuli and task complexity but differed in action outcome. During JA, participants achieved a collective goal by acting jointly with their partner, while during PA, they pursued an individual goal, acting merely side by side with their partner.

**Figure 2:**
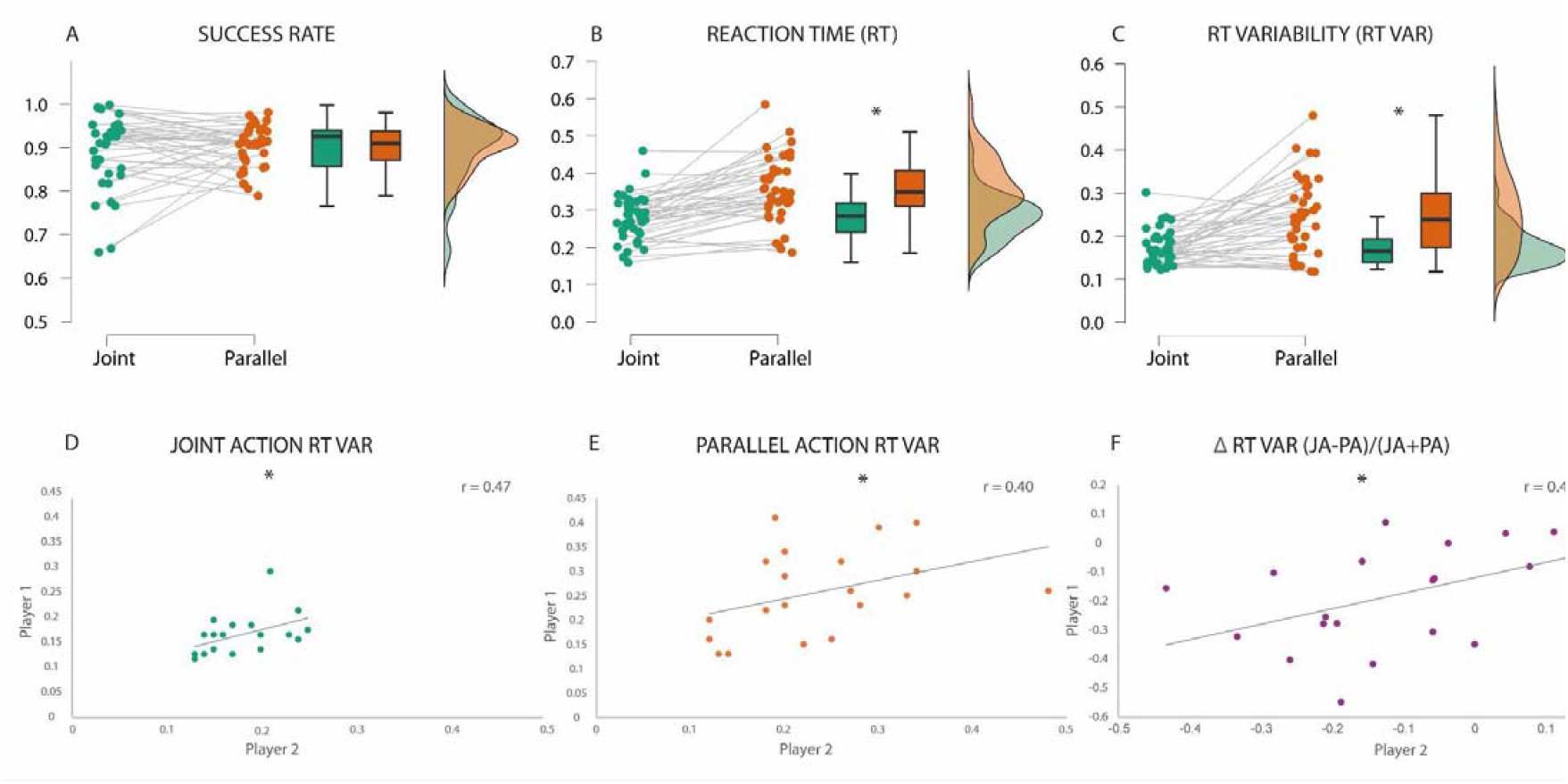
Behavioral results. The top panels show the significant differences between JA and PA conditions in (A) success rate, (B) reaction times, and (C) reaction time variability. Each panel shows individual participants’ performance on the left, boxplots in the middle, and density distribution on the right. The bottom panels show significant correlations in reaction time variability across pairs of players. Here, each dot represents the performance of the two players of the couple. Specifically, (D) shows the correlation in the JA condition, (E) shows the correlation in the PA condition, and (F) shows the correlation of ΔRTvar.

**Figure 3:**
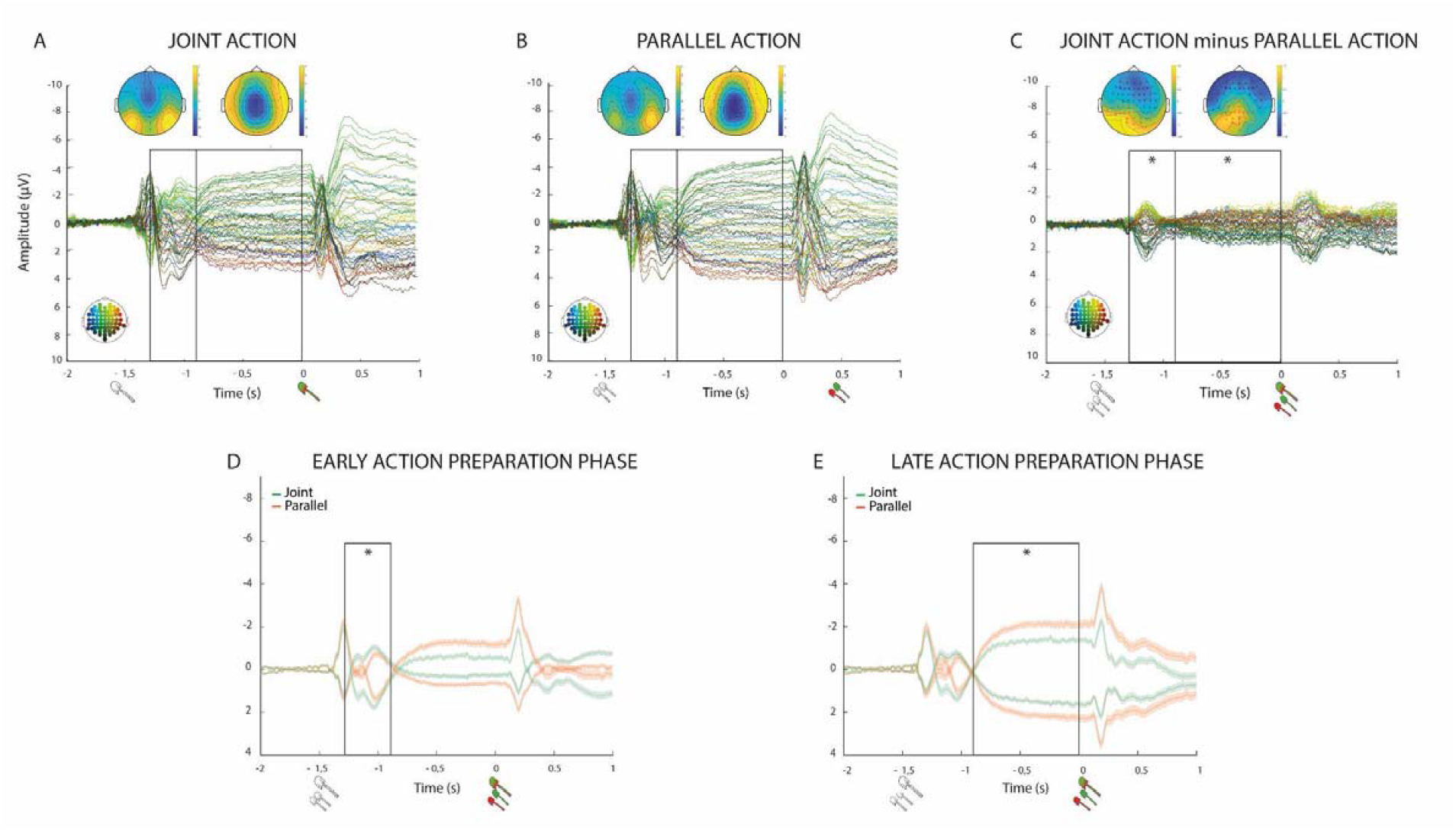
Difference in ERPs between JA and PA conditions (all ps<0.005). (A) and (B) show the butterfly plots of ERP measured in the JA and PA conditions, respectively. Each line represents the signal recorded in an electrode, as indicated by the color code in the head at the bottom of the graph. Topographies above the butterfly plots represent the average topography in the time windows indicated by the black rectangles. (C) shows the butterfly plot of the difference of ERPs (JA-PA). Black rectangles indicate the time windows of the significant clusters. Topographies above the rectangles indicate the spatial distribution of the clusters; red asterisks represent positive clusters, and black asterisks represent negative clusters. The average of the signals from the electrodes of each cluster is displayed in the bottom panel: (D) depicts the average signal of the positive and negative clusters for the early window; (E) depicts the average signal of the two clusters in the late time window. White shovels represent the time of the cue signal, and colored shovels represent the time of the go signal.

**Figure 4.**
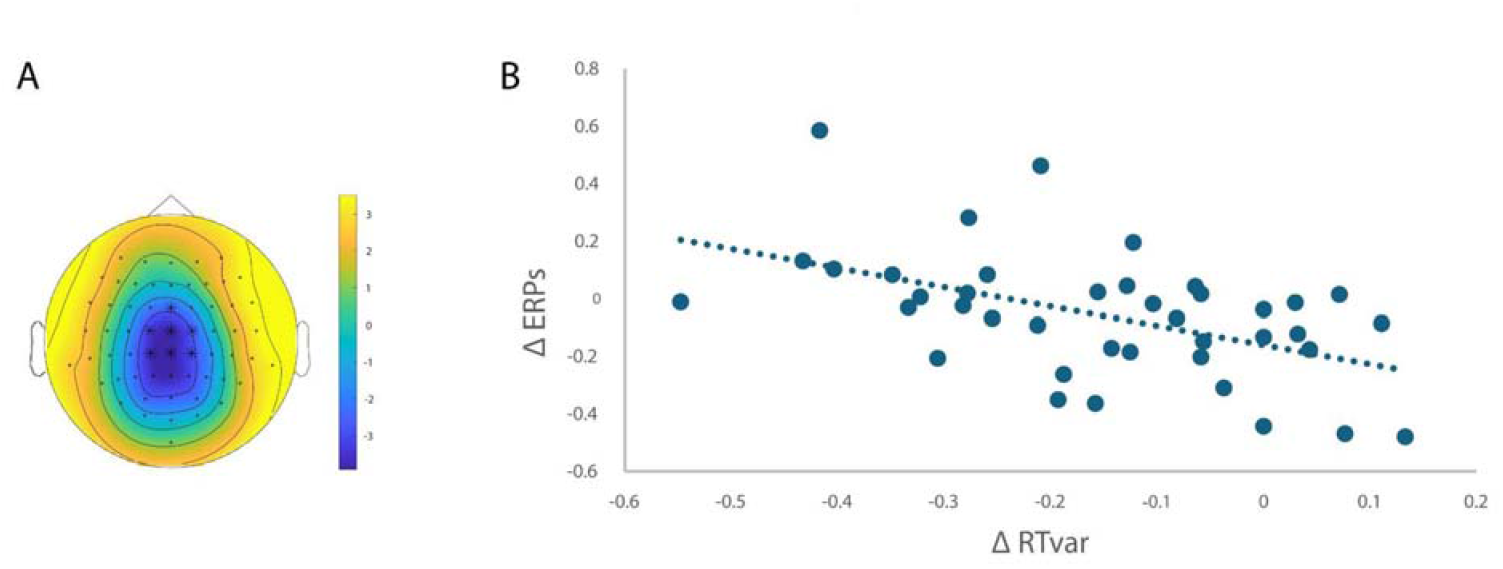
Correlation between ΔERPs and ΔRTvar. A significant difference was found in the cluster-based correlation analyses (p = 0.036). (A)The topographic plot shows the electrodes included in the significant cluster (highlighted as black dots), superimposed on the average topography of the ERPs in JA and PA within the cluster time window. (B) Graphs show the correlation between ΔERPs in the cluster and the ΔRTvar. For display purposes, we calculated the cluster’s mean value for each participant and plotted the correlation between these values and ΔRTvar.

Participants completed eight blocks of 48 trials each (384 total trials, 192 per condition), with 12 repetitions for each of the four target directions (up-left, up-right, down-left, down-right). JA and PA blocks alternated, with starting conditions counterbalanced across dyads. Before the experiment, each dyad completed two training blocks of 16 trials each (4 directions × 4 repetitions), following their assigned experimental order (JA-PA or PA-JA). Training trials were excluded from the analysis. All procedures were administered via Psychtoolbox-3 in MATLAB 2021b.

### Behavioral data analysis

Trials were included in the analyses only if they presented a clear post-go movement onset, defined as the first velocity peak after the go-signal exceeding two mean absolute deviations above the median velocity in the action preparation phase. In this way, we excluded both trials with no responses and trials in which excessive movement activity was detected during the preparation phase.

On the remaining trials, we calculated the Success rate (SR) as the percentage of trials in which participants successfully reached the igloo, the Reaction Times (RTs) as the interval between go-signal and movement onset, and the Reaction time variability (RTvar) as the mean standard deviation of RTs across trials. These variables were analyzed separately in 2×2 mixed ANOVAs, with the within-subject factor Condition (JA, PA) and the between-subject factor Player (Red, Green). The latter factor was considered to ensure that the players’ positions did not affect their performance.

Moreover, we conducted an exploratory analysis of dyadic performance to examine behavioral alignment within pairs of participants. We analyzed the linear relationship between players’ RT and RTvar by calculating correlations between paired participants across all 20 dyads. We specifically tested for positive correlations, indicating alignment, as negative correlations would indicate a lack thereof. To quantify how changes in behavioral performance between JA and PA conditions correlated within dyads, we computed the normalized difference ((JA-PA)/(JA+PA)) for both mean reaction time (ΔRT) and reaction time variability (ΔRTvar) for each participant. Then, we separately calculated correlations of ΔRT and ΔRTvar between paired participants across all 20 dyads.

Analyses were run using Jamovi 2.3.24 and JASP 0.18.3.0.

### ERP analysis

#### ERP data preprocessing

Data were preprocessed using EEGLAB (version 2023.1), implemented in MATLAB 2023 b. First, continuous data were epoched into 8-second segments (−2000 to 6000 ms relative to the instruction cue), baseline-corrected using the pre-stimulus period, downsampled to 600 Hz, linearly detrended, and low-pass filtered at 30 Hz (using a FIR filter with default settings). After visual inspection, bad channels were interpolated (mean ± sd: 0.9 ± 0.8 channels), and data were average-referenced. Trials with excessive artifacts were removed upon visual inspection, followed by ICA decomposition (RUNICA with default settings). After removing components that had been visually identified as ocular artifacts (mean ± sd: 3.3 ± 0.9 components), trials were re-epoched in 2500 ms segments, starting from 500 ms before the visual cue and ending 500 ms after the go-signal. These epochs were baseline-corrected to 500 ms before the visual cue. A second trial rejection was then performed to eliminate remaining bad trials and trials in which no clear movement onset had been detected (mean number of trials included ± sd: 112.25 ± 31 for JA and 131 ± 24 for PA). Finally, the signal was averaged across trials.

#### Cluster-based permutation analysis

To investigate the differences in ERPs between JA and PA conditions during the entire action preparation phase, we performed a cluster-based analysis as implemented in FieldTrip (Maris & Oostenveld, 2007; Oostenveld et al., 2011). First, we conducted dependent sample t-tests to examine differences in ERP amplitudes between the JA and PA conditions. These comparisons were performed across all electrodes and time samples within the 1500 ms interval between the visual and go cues. Clusters were formed by two-tailed t-values with a p-value < 0.05, considering adjacent time samples and neighboring electrodes (n ≥ 2), as defined with default triangulation parameters. To assess significance, the summed t-value of each cluster (cluster statistic =maxsum) was compared against the distribution of clusters obtained through permutation testing. Using the Monte Carlo method, ERPs were randomly assigned to either the JA or PA condition 1000 times. The permutation test was two-tailed, and clusters were deemed significant if fewer than 5% of the summed t-values from the permutations exceeded the original cluster t-value (p-value < 0.05).

#### Correlations between task-related changes in ERPs and behavior

We finally explored the potential correlations between ERPs and behavioral outcomes. In doing this, we calculated the point-by-point normalized difference between JA and PA ((JA-PA)/(JA+PA)) of the ERP amplitudes during action preparation (ΔERP). With separate cluster-based correlations, we tested its association with partner and self-behavior changes, specifically ΔRT and ΔRTvar. We ran correlations with ΔERPs across all electrodes and time samples for each behavioral variable within the 1500 ms interval between the instruction and go cues. Then, clusters were formed using two-tailed t-values with a p-value < 0.05, considering channel neighbors (n ≥ 2) defined by a triangulation-based layout and adjacent time points. The statistical significance of clusters was determined via a Monte Carlo method with 1000 randomizations, using summed t-values as the cluster statistic (=maxsum). Clusters were considered significant for p<0.05.

Considering that this normalization is impacted when conditions’ values are close to zero, leading to issues with dividing by small numbers, we controlled the percentage of denominator values smaller than 0.1 in the significant clusters.

## RESULTS

### Behavioral Results

We found no significant differences in SR between JA and PA conditions (F_(1, 38_) = 0.136, p = 0.577, pη^2^ = 0.008; JA: M = 0.897, SD = 0.081; PA: M = 0.903, SD = 0.048). The factor player (Red/Green) showed no main effect (p = 0.356) or interaction with Condition (p = .503), indicating that seating position did not affect performance.

Mean RTs were significantly higher in PA than in JA condition (F(_1, 38_) = 42.924, p < 0.001, pη^2^ = 0.530; JA: M = 0.329, SD = 0.062; PA: M = 0.406, SD = 0.089). The factor player showed no main effect or interaction with Condition for either mean RTs (main effect: p = 0.496; interaction: p = 0.949).

Similar to RTs, RTvar was greater in PA than in JA condition (F(_1, 38_) = 29.801, p < 0.001, pη^2^ = 0.440; JA: M = 0.173, SD = 0.040; PA: M = 0.247, SD = 0.089), with no effect of the player factor (main effect: p = 0.752; interaction: p = 0.295).

RTs were correlated between paired participants across all 20 dyads in the JA (r = 0.43, p = 0.03) and PA (r = 0.42, p = 0.03) conditions. This result indicated that participants adjusted their response times (RTs) to match those of their partner. Specifically, participants tended to respond slowly when their partner’s responses were slow; conversely, when their partner’s responses were fast, they tended to respond quickly.

RTvars were also positively correlated in JA (r = 0.47, p = 0.02) and PA (r = 0.40, p = 0.04), indicating that variability in initiating movements was adjusted to match the partner’s response pattern.

Finally, behavioral performance changes between the JA and PA conditions correlated within dyads. Precisely, changes in ΔRTvar for the two partners matched, so greater RTvar in PA than in JA in one player tended to correspond to greater RTvar in PA than in JA in their partner (r = 0.46, p = 0.02). No significant correlation was found for ΔRT (r = 0.19, p = 0.21).

### ERP Results

JA and PA conditions differed significantly around two time windows, with a reversed pattern. A first, early difference revealed a higher ERP amplitude in the JA compared to the PA condition. This was evident as enhanced negativity in a fronto-central cluster of electrodes (p = 0.002; time: from -1.250 to -0.882 ms) and enhanced positivity in JA compared to PA in an occipito-parietal cluster (p = 0.004; time: from -1.261 to -0.898 ms). A second, late difference revealed a smaller ERP amplitude in the JA compared to the PA condition. Accordingly, we found significantly reduced negativity in central-parietal channels (p = 0.002; time: from -0.883 to 0 ms) and reduced positivity in fronto-lateral channels (p = 0.002; time: from -0.820 to 0 ms) during JA compared to the PA condition. Although the time intervals for significant clusters should not be considered as exact latencies in cluster-based analyses (Sassenhagen and Draschkow, 2019), these results clearly show a reverse pattern over two distinct time windows.

### Correlations between task-related changes in ERP activity and behavior

Correlations between ERPs and behavioral outcomes showed no significant relationship between ΔERPs and changes in self-behavioral performance, ΔRT, and ΔRTvar (p>0.05). Interestingly, we found a cluster of significant correlation between ΔERPs and partner’s ΔRTvar (clusters: p= 0.036 from -0.448 to -0.402 ms) and two clusters suggesting a trend in the same direction (p= 0.068 from -0.053 to 0.018 ms; p = 0.082 from -0.387 to -0.350 ms). All these clusters occurred in the late part of the preparation phase. They included frontocentral channels, indicating that a stronger increase in negative ERP amplitude in PA compared to JA was associated with a greater increase in variability of movement initiation in PA than in JA. The control analyses supported that small denominators do not drive these results, as less than 0.1% of denominators were smaller than 0.1 µV in each cluster. No significant correlation was found for ΔRT.

## DISCUSSION

The main aim of our paper was to investigate behavioral patterns and neural dynamics that distinguish joint from parallel action preparation. We analyzed mutual alignment measured by differences in reaction time (RT) and reaction time variability (RTvar) and compared event-related potentials (ERPs) simultaneously recorded from pairs of participants while they prepared to play a video game in which they had to transport a shovel *jointly* or in *parallel*.

We found that the success rate did not differ across conditions, indicating that the tasks were well-matched for task complexity. Participants adjusted their RTs and RTvars to match their partner’s response pattern in both the Joint and Parallel conditions, with reduced overall variability observed in the Joint condition. The findings indicate that participants considered the partner’s actions and coordinated in both conditions, with mutual adjustment generally smoother when they acted jointly.

ERP analysis revealed an intriguing swap in differences in ERP amplitudes between joint and parallel action preparation. Indeed, we found two distinct time windows with reversed patterns. A first, early time window (approximately from -1.250 ms to -880 ms) revealed a higher ERP amplitude in the Joint than in the Parallel condition, characterized by enhanced fronto-central negativity and occipito-parietal positivity. In contrast, the second, late time window (from -880 ms to 0) showed a smaller ERP amplitude in the Joint condition than in the Parallel condition, with reduced central-parietal negativity and fronto-lateral positivity.

Strikingly, the differences in ERP between the Joint and Parallel conditions significantly correlated with the differences in the partner’s RTvar in frontocentral channels during the late action preparation phase (approximately -400 ms). Notably, enhanced condition-related ERP negativity corresponded to a greater increase in partner movement initiation variability in the Parallel condition compared to the Joint condition. No correlation with self-behavioral changes was found.

These results showed the typical ERP components reported in CNV paradigms, including evoked responses to the stimuli and the CNV in the interval between them (Walter et al., 1964). After its discovery, CNV was refined in components: an early orienting wave (O-wave), occurring around 450-650 ms after the cue and larger at fronto-central sites, and a later expectancy wave (E-wave), larger at centro-parietal sites (Loveless and Sanford, 1974). The first component is linked to cognitive processing and the properties of the first stimulus, and the second component is associated with expectancy and motor preparation (Weerts and Lang, 1973; Rohrbaugh and Gaillard, 1983). Although these two components have been fully distinguished when the interval between stimuli is longer than in our study, they are still observable in our data, suggesting that both the early orienting response and the later expectancy-related activity contributed to the CNV, even within the shorter interstimulus interval.

The swap we found corresponds well to the functional distinction of the O-wave and the E-wave components of the CNV. The first difference occurred in the time window of the O-wave and consisted of a higher amplitude for the Joint than the Parallel condition. The second difference, in which the amplitude is higher for the Parallel than the Joint Action condition, was in the time range of the E-wave. Therefore, the early effect likely relies more on the O-wave while the later effect relies more on the E-wave.

A characterization of the distinctive functions associated with the O-wave of the CNV is lacking. Nevertheless, there is evidence that it may primarily reflect cognitive processes providing a general preparedness to respond, such as stimulus processing, orientation, and anticipation in response to warning signal (Brunia and van Boxtel, 2001; Gómez et al., 2019). The O-wave is modulated by a broad spectrum of factors (Rohrbaugh and Gaillard, 1983), such as stimulus features, cue saliency, and reward. Interestingly, specific motor preparation processes have been shown to already occur at the time of the O-wave (Bender et al., 2004). Therefore, it may be possible that this early response represents the integration of several task-related aspects, linking the processing of the cue with the representation of the goal. This could explain why the early response is higher in the Joint condition, when participants had to represent a collective goal, than in the Parallel condition, where collective goal representation was absent. The fact that the stimuli presented in both conditions were virtually identical corroborated this explanation.

The reduction of the E-wave in the Parallel compared to the Joint condition likely reflects motor preparation changes rather than temporal expectation. Indeed, the late CNV component related to temporal expectation is reduced when there is more temporal uncertainty on the go stimulus (Breska and Deouell, 2014; Praamstra et al., 2006; Trillenberg et al., 2000; Duma et al., 2020; Mento et al., 2013). However, the go-signal did not differ in temporal uncertainty across the conditions. Furthermore, the CNV was reduced in the Joint condition, which was characterized by a lower temporal response variability. This suggests that our results are unlikely to be explained mainly by temporal expectancy. Instead, a more likely explanation is that the reduction of late CNV in the Joint condition is related to the modulation of motor preparation processes.

This explanation fits well with the significant association between the change in CNV amplitude and the variability of the partner’s movement, which was reduced in the Joint compared to the Parallel condition. The less variable the movements are, the more predictable they are (Vesper et al., 2013; Sabu et al., 2020; also, Keller et al., 2007). The higher predictability of partner movements might, therefore, reduce demands on participants’ motor planning in the Joint compared to the Parallel condition, as the number of alternative movements that may have to be executed while coordinating with the confederate is limited, and movement parameters tend to be fixed across task repetitions. This is also in line with studies on CNV with ambiguous pre-cues (Jentzsch et al., 2004) and on readiness potential during action selection (Dirnberger et al., 1998; Praamstra et al., 1995; Touge et al., 1995), which all exhibited a reduction of the potential when movement variability was limited.

If our explanation is correct, the dynamic swap between early and late CNV components reveals fundamentally distinct processes between joint and parallel action preparation. The cognitive demands of representing collective goals may initially enhance neural activity associated with the early stages of action preparation, but these costs are offset later by enhanced mutual predictability, which facilitates action preparation. Participants considered partners’ actions when acting in parallel, which enabled successful coordination. However, joint action involves more than considering others’ actions (Bratman 2014; Ludwig 2016). Agents plan their own and others’ actions collectively toward a shared goal. When these representations sufficiently match, as happened with participant dyads, action initiation is facilitated, increasing the likelihood of achieving the collective goal (Butterfill & Sinigaglia, 2023).

Our findings appear to conflict with previous studies (Kourtis et al., 2014, 2019), which reported a greater late CNV amplitude when agents act together compared to when they act alone, attributing this to the representation of the other agent’s actions. However, the conflict is more apparent than real. Indeed, performing a task with someone else is usually more complex than performing the same task alone, and evidence shows that late CNV increases for more complex tasks (Kranczioch, Mathews, Dean, & Sterr, 2010; Cui et al., 2000). Our comparison of complexity-matched Joint and Parallel tasks revealed reduced late CNV amplitudes in the Joint condition, correlating with increased predictability of the partner when shared goals were involved. Participants needed to consider their partners’ movements in both tasks, yet distinctive late CNV patterns emerged. This suggests late CNV variations reflect others’ action predictability rather than merely representing their actions (which occurs in both conditions). This interpretation also reconciles the increase in CNV when comparing solo action with joint action, as actions are more predictable when acting alone than when acting with others.

Finally, our results are consistent with recent studies that demonstrate planning and performing complementary actions create more significant interference when acting in parallel than when acting jointly. Formica & Brass (2024) found that drawing incongruent shapes (e.g., circle and diamond) disrupted participants’ trajectories more in the Parallel than in the Joint condition (see also Sartori & Betti, 2015, and Clarke et al., 2019). Interestingly, their EEG analysis showed that the incongruent movements were more discriminable in the Parallel than in the Joint condition. As the authors suggest, this may result from forming an intertwined representation when acting jointly but not in parallel. Our results extend this suggestion, revealing that sharing a collective goal representation enhances action predictability by reducing interindividual variability at both behavioral and electrophysiological levels. Moreover, these findings may have important clinical implications, particularly for psychiatric conditions characterized by social interaction deficits (Schilbach, 2016), such as schizophrenia, where CNV has emerged as a promising neurophysiological marker (Akgül et al., 2023).

Although further research is needed, our study provides a clear insight into the behavioral and neurophysiological processes that distinguish joint from parallel actions. Acting jointly entails a dynamic trade-off: the initial cognitive investment in representing collective goals yields benefits during late-stage action preparation through enhanced mutual predictability.

## Acknowledgments

This article was supported by the Department of Philosophy ‘Piero Martinetti’ of the University of Milan with the Project “Departments of Excellence 2018-2022” and “Departments of Excellence 2023-2027” awarded by the Italian Ministry of Education, University and Research (MIUR) (to MF, GB, and CS), the PRIN 2017 grant “The cognitive neuroscience of interpersonal coordination and cooperation: a motor approach in humans and non-human primates” (201794KEER; to ABM, and CS), the PRIN 2022 grant “The extended hand: psychophysical and neural foundations of a robotic supernumerary finger’s use for grasping augmentation or recovery” (2022J72LFW_002 to CS), the PRIN 2022 grant “Motor resonance during action planning and social interactions: from single neurons to brain circuits” (2022SP5K99_002 to GB) and Italian Ministry of Health (‘Ricerca Corrente’ to AZ and MB).

## REFERENCES

1. Akgül Ö, Fide E, Özel F, Alptekin K, Bora E, Akdede BB, Yener G (2024) Early and late contingent negative variation (CNV) reflect different aspects of deficits in schizophrenia. Eur J Neurosci 59(11):2875–2889.

2. Atmaca S, Sebanz N, Knoblich G (2011) The joint flanker effect: sharing tasks with real and imagined co-actors. Exp Brain Res 211:371–385.

3. Atmaca S, Sebanz N, Prinz W, Knoblich G (2008) Action co-representation: the joint SNARC effect. Soc Neurosci 3:410–420.

4. Barchiesi G, Zazio A, Marcantoni E, Bulgari M, di San Pietro CB, Sinigaglia C, Bortoletto M (2022) Sharing motor plans while acting jointly: A TMS study. Cortex 151:224–39.

5. Bender S, Resch F, Weisbrod M, Oelkers-Ax R (2004) Specific task anticipation versus unspecific orienting reaction during early contingent negative variation. Neurophysiol Clin 115(8); 1836–1845.

6. Bigand F, Bianco R, Abalde SF, Novembre G (2024) The geometry of interpersonal synchrony in human dance. Curr Biol 34(13):3011–3019

7. Bratman M (1993) Shared intention. Ethics 104:97–11.

8. Bratman M (2014) Shared Agency. Oxford: Oxford University Press.

9. Breska A, Deouell LY (2014) Automatic bias of temporal expectations following temporally regular input independently of high-level temporal expectation. J Cogn Neurosci 26(7);1555–1571.

10. Brownell CA (2013) Early development of prosocial behaviour. Current perspectives. Infancy 18 (1): 1–9.

11. Brunia CH, van Boxtel, GJ (2001) Wait and see. Int J Psychophysiol 43(1), 59–75.

12. Butterfill SA, Sinigaglia C (2023) Towards a mechanistically neutral account of acting jointly: The notion of a collective goal. Mind 132(525):1–29.

13. Carpenter M (2009) Just How Joint Is Joint Action in Infancy? Top Cogn Sci 1:380–392.

14. Clarke S, McEllin L, Francová A, Székely M, Butterfill SA, Michael J (2019) Joint action goals reduce visuomotor interference effects from a partner’s incongruent actions. Sci Rep 9(1):15414.

15. Cui RQ, Egkher A, Huter D, Lang W, Lindinger G, Deecke L (2000) High resolution spatiotemporal analysis of the contingent negative variation in simple or complex motor tasks and a non-motor task. Neurophysiol Clin 111(10):1847–1859.

16. D’Ausilio A, Novembre G, Fadiga L, Keller PE (2015) What can music tell us about social interaction?. Trends Cogn Sci 19(3):111–114.

17. della Gatta F, Garbarini F, Rabuffetti M, Viganò L, Butterfill SA, Sinigaglia C (2017) Drawn together: When motor representations ground joint actions. Cognition 165:53–60

18. Dirnberger G, Fickel U, Lindinger G, Lang W, Jahanshahi M (1998) The mode of movement selection. Exp Brain Res 120(2):263–272.

19. Duma G M, Granziol U, Mento G (2020) Should I stay or should I go? How local-global implicit temporal expectancy shapes proactive motor control: An hdEEG study. NeuroImage 220:117071.

20. Ferrari-Toniolo S, Visco-Comandini F, Battaglia-Mayer A (2019) Two Brains in Action: Joint-Action Coding in the Primate Frontal Cortex. J Neurosci. 39(18):3514–3528.

21. Formica S, Brass M (2024) Coordinated social interactions are supported by integrated neural representations. Soc Cogn Affect Neurosci 19(1):nsae089.

22. Gilbert M (1990) Walking together: A paradigmatic social phenomenon. Midwest Studies in Philosophy 15:1–14.

23. Gilbert M (2013) Joint Commitment: How We Make the Social World. Oxford: Oxford University Press.

24. Gómez CM, Arjona A, Donnarumma F, Maisto D, Rodríguez-Martínez EI, Pezzulo G (2019) Tracking the time course of bayesian inference with event-related potentials: A study using the central cue Posner paradigm. Front Psychol 10:1424.

25. Issartel J, Marin L, Cadopi M (2007) Unintended interpersonal co-ordination: ‘Can we march to the beat of our own drum?’ Neurosci Lett 411 (3):174–179.

26. Jentzsch I, Leuthold H, Ridderinkhof KR (2004) Beneficial effects of ambiguous precues: Parallel motor preparation or reduced premotoric processing time? Psychophysiology 41(2):231–244.

27. Keller PE, Knoblich G, Repp BH (2007) Pianists duet better when they play with themselves: on the possible role of action simulation in synchronization. Conscious Cogn. 16(1):102–11.

28. Keller PE, Novembre G, Hove MJ (2014) Rhythm in joint action: psychological and neurophysiological mechanisms for real-time interpersonal coordination. Philos Trans R Soc Lond B Biol Sci 369(1658):20130394.

29. Konvalinka I, Vuust P, Roepstorff A, Frith CD (2010) Follow you, follow me: continuous mutual prediction and adaptation in joint tapping. Quar J Exp Psychol 63(11):2220–230.

30. Kourtis D, Knoblich G, Wozniak M, Sebanz N (2014) Attention, allocation and task representation during joint action planning. J Cogn Neurosci 26:2275–2286.

31. Kourtis D, Woźniak M, Sebanz N, Knoblich G. (2019) Evidence for we-representations during joint action planning. Neuropsychologia 131:73–83.

32. Kranczioch C, Mathews S, Dean P, Sterr A (2010) Task complexity differentially affects executed and imagined movement preparation: evidence from movement-related potentials. PloSOne 5(2);e9284.

33. Lacal I, Babicola L, Caminiti R, Ferrari-Toniolo S, Schito A, Nalbant LE, Gupta RK, Battaglia-Mayer A (2022) Evidence for a we-representation in monkeys when acting together. Cortex 149:123–136.

34. Lelonkiewicz JR, Gambi C (2017) Spontaneous adaptation explains why people act faster when being imitated. Psychon Bull Rev 24:842–488.

35. Loehr JD, Kourtis D, Vesper C, Sebanz N, Knoblich G (2013) Monitoring individual and joint action outcomes in duet music performance. J Cogn Neurosci 25:1049– 1061.

36. Loveless NE, Sanford A J (1974) Slow potential correlates of preparatory set. Biol Psychol 1(4):303–314.

37. Ludwig K (2016) From Individual to Plural Agency. Collective Action: Volume 1. Oxford: Oxford University Press

38. Maris E, Oostenveld R (2007) Nonparametric statistical testing of EEG- and MEG-data. J Neurosci Meth 164(1):177–190.

39. Mento G, Tarantino V, Sarlo M, Bisiacchi PS (2013) Automatic temporal expectancy: a high-density event-related potential study. PloSOne 8(5);e62896.

40. Meyer M, van der Wel RP, Hunnius S (2013) Higher-order action planning for individual and joint object manipulations. Exp Brain Res 225:579–588.

41. Nessler JA, Gilliland SJ (2009) Interpersonal synchronization during side by side treadmill walking is influenced by leg length differential and altered sensory feedback. Hum Mov Sci 28:772–785.

42. Novembre G, Ticini LF, Schutz-Bosbach S, Keller PE (2014) Motor simulation and the coordination of self and other in real-time joint action. Soc Cogn Affect Neurosci 9:1062–1068.

43. Nowicki L, Prinz W, Grosjean M, Repp BH, Keller PE. (2013) Mutual adaptive timing in interpersonal action coordination. Psychomusicology 23(1):6.

44. Oostenveld R, Fries P, Maris E, Schoffelen JM (2011) FieldTrip: open source software for advanced analysis of MEG, EEG, and invasive electrophysiological data. Comput Intell Neurosci. 9 pages.

45. Pezzulo G, Donnarumma F, Ferrari-Toniolo S, Cisek P, Battaglia-Mayer A (2022) Shared population-level dynamics in monkey premotor cortex during solo action, joint action and action observation. Progr Neurobiol, 210:102214.

46. Pezzulo G, Iodice P, Donnarumma F, Dindo H, Knoblich G (2017) Avoiding accidents at the champagne reception: A study of joint lifting and balancing. Psychol Sci 28(3):338–345.

47. Praamstra P, Kourtis D, Hoi FK, Oostenveld R (2006) Neurophysiology of implicit timing in serial choice reaction-time performance. J Neurosci 26(20):5448–5455.

48. Praamstra P, Stegeman DF, Horstink MW, Brunia CH, Cools AR (1995) Movement-related potentials preceding voluntary movement are modulated by the mode of movement selection. Exp Brain Research 103(3):429–439.

49. Repp BH, Su YH (2012) Sensorimotor synchronization: a review of recent research (2006–2012). Psychon Bull Rev 20:403–52.

50. Richardson MJ, Marsh KL, Isenhower RW, Goodman JR, Schmidt R (2007) Rocking together: Dynamics of intentional and unintentional interpersonal coordination. Hum Mov Sci 26:867–891.

51. Rohrbaugh JW, Gaillard AW (1983) 13 Sensory and motor aspects of the contingent negative variation. Adv Psychol 10(C):269–310.

52. Sabu S, Curioni A, Vesper C, Sebanz N, Knoblich G (2020) How does a partner’s motor variability affect joint action? PloSOne 15(10):e0241417.

53. Sacheli LM, Arcangeli E, Paulesu E (2018) Evidence for a dyadic motor plan in joint action. Scie Rep 8(1):5027.

54. Sartori L, Betti S (2015) Complementary actions. Front Psychol. 6:557.

55. Sassenhagen J, Draschkow D (2019) Cluster-based permutation tests of MEG/EEG data do not establish significance of effect latency or location. Psychophysiology 56(6):e13335.

56. Satta E, Ferrari-Toniolo S, Visco-Comandini F, Caminiti R, Battaglia-Mayer A (2017) Development of motor coordination during joint action in mid-childhood. Neuropsychologia 105:111–122.

57. Schilbach L (2016) Towards a second-person neuropsychiatry. Philos Trans R Soc Lond B Biol Sci 371(1686):20150081.

58. Schmitz L, Vesper C, Sebanz N, Knoblich G (2018) Co-actors represent the order of each other’s actions. Cognition 181:65–79.

59. Schmitz L, Vesper C, Sebanz N, Knoblich G (2017) Co-representation of others’ task constraints in joint action. J Exp Psychol: Hum Percept Perform 43(8):1480.

60. Searle J (1990) Collective intentions and actions. In: Intentions in Communication (Cohen P, Morgan J, Pollack ME eds.) Cambridge, Mass.: MIT Press.

61. Sebanz N, Knoblich G (2021) Progress in joint-action research. Curr Dir Psychol Sci 30(2):138–143.

62. Sebanz N, Knoblich G, Prinz W (2003) Representing others’ actions: just like one’s own? Cognition 88:B11–B21.

63. Tomasello M, Carpenter, M, Call J, Behne T, Moll H (2005) Understanding and sharing intentions: The origins of cultural cognition. Behav Brain Sci 28:675–735.

64. Touge T, Werhahn KJ, Rothwell JC, Marsden CD (1995) Movement-related cortical potentials preceding repetitive and random-choice hand movements in parkinson’s disease. Ann Neurol 37(6); 791–799.

65. Trillenberg P, Verleger R, Wascher E, Wauschkuhn B, Wessel K (2000) CNV and temporal uncertainty with “ageing” and “non-ageing” S1-S2 intervals. Neurophysiol Clin 111(7):1216–1226.

66. Ulzen NR van, Lamoth CJ, Daffertshofer A, Semin GR, Beek PJ (2008) Characteristics of instructed and uninstructed interpersonal coordination while walking side-by-side. Neurosci Lett 432 (2):88–93.

67. Varlet M, Bucci C, Richardson MJ, Schmidt RC (2015) Informational constraints on spontaneous visuomotor entrainment. Hum Mov Sci 41:265–281.

68. Vesper C, van der Wel RP, Knoblich G, Sebanz N (2013) Are you ready to jump? Predictive mechanisms in interpersonal coordination. J Exp Psychol: Hum Percept Perform 39 (1):48–61.

69. Walter WG, Cooper R, Aldridge VJ, McCallum WC, Winter AL (1964) Contingent negative variation: An electric sign of sensorimotor association and expectancy in the human brain. Nature 203:380–384.

70. Weerts TC, Lang PJ (1973) The effects of eye fixation and stimulus and response location on the Contingent Negative Variation (CNV). Biol Psychol 1(1):1–19.

